# Principles for Optimal Window Size Selection for Infant and Adult EEG Connectivity Analysis

**DOI:** 10.1101/2023.05.31.542996

**Authors:** Lorena Santamaria, Andres Canales-Johnson, Valdas Noreika, Victorial Leong

## Abstract

Neural connectivity analysis is often performed on continuous data that has been discretized into temporal windows of a fixed length. However, the selection of an optimal window length is non-trivial, and depends on the properties of the connectivity metric being used as well as the effects of interest within the data (e.g. developmental or inter-brain effects). A systematic investigation of these factors, and objective criteria for window size selection are currently missing in the literature, particularly in regard to pediatric datasets. Here, we provide a principled examination of the effect of window size on optimization of signal to noise ratio for linear and non-linear EEG connectivity, as applied to infant, adult and dyadic (infant-adult) datasets. We employed a linear weighted phase lag index (wPLI), and a nonlinear weighted symbolic mutual information (wSMI) metric to assess brain connectivity for each dataset. Our results showed a clear polar dissociation between linear and non-linear metrics, as well as between infant and adult datasets in optimal window size. Further, optimal dyadic (infant-adult) window size settings defaulted to one or the partner rather than reflecting an intermediate compromise. Given the specificity of these results (i.e. there was no single window size that was optimal for all contrasts), we conclude that a formal analysis of optimal window size may be useful prior to conducting any new connectivity analysis. Here, we recommend guiding principles, performance metrics and decision criteria for optimal and unbiased window size selection.

## I. Introduction

Electroencephalography (EEG) measures electrical potentials recorded from the surface of the scalp. These are the product of complex neural interactions combining oscillatory and non-oscillatory dynamics [1]. Whilst oscillatory dynamics are characterized by distinctive peaks in the power spectrum (e.g., alpha 10 Hz), non-oscillatory dynamics reflect neural activity that does not correspond to a specific rhythm or temporal pulse. Yet both forms of neural activity are known to support neural connectivity (and communication) between brain regions [1]. A lack of rhythmicity complicates the quantification of non-oscillatory interactions using traditional brain connectivity methods that are based on spectral decomposition, such as Coherence, Granger Causality, or the weighted phase lag index (wPLI), all of which are linear methods. Yet non-linear interactions are known to be essential for pattern extraction in neural networks (e.g. object or speech recognition) and may lead to signal amplification and broadcasting via non-linear recurrent dynamics (“ignition”) [2].

One method that is able to capture non-oscillatory interactions is the wSMI (weighted symbolic mutual information). This non-linear metric quantifies the extent to which the temporal structure of one signal predicts the temporal structure of a second signal. Crucially, wSMI detects differences between perceptual states that are not picked up by traditional linear measures [3]. For instance, wSMI outperforms linear connectivity metrics (e.g. wPLI) when separating pathological states of consciousness in vegetative and minimally conscious patients [4]. Further, during active tasks, fronto-parietal wSMI (but not wPLI) distinguishes between perceptual interpretations of a bistable stimulus in human participants [5].

A further consideration is that the relative contribution and importance of linear versus non-linear brain dynamics may change during the course of human development. During early life, infant neural dynamics are dominated by low-frequency spectral power, with highest power in the infant theta and alpha ranges of the EEG signal (3-9 Hz). However, it is not known whether non-linear brain dynamics already exist at an early age, and what is their relative contribution toward cognitive processes. One possibility is that the temporal structure of neural interactions may change during development, moving progressively away from dominant oscillatory dynamics toward non-oscillatory dynamics. This would suggest that linear connectivity metrics such as the wPLI may be more effective in capturing early neural connectivity patterns than non-linear metrics such as wSMI. During adulthood, the converse may be true. Accordingly, it is of interest to study whether developmental differences in linear versus non-linear dynamics exist, and what are their consequences for developing cognition.

Both linear and non-linear patterns of neural activity are often studied using graph network measures. Using this approach, neural connectivity is represented as a graph whereby nodes represent brain regions and edges denote connections through which information is passed between brain areas. Measures of functional connectivity (FC), usually represented in matrix format, have been extensively used in neuroscience research to elucidate principles of neural function [6]. Further, studies have demonstrated that functional connectivity does not remain temporally static, but rather changes dynamically over time [7]. One common method used to study these temporal dynamics is the sliding window approach [8]. This has been successfully used with adult data, in both healthy [9], and clinical populations [10], as well as in infant cohorts [11]. However, the selection of an optimal window size is not a trivial matter. There exists a trade-off between keeping the window small enough to capture (fast) temporal transitions yet large enough to yield stable results [12]. The picture is further complicated by whether linear or non-linear neural dynamics are functionally more relevant. Non-linear dynamics may require longer time windows in order to increase the probability of observing distinct temporal patterns jointly occurring over time. Finally, there are developmental considerations as to the most appropriate window size for pediatric versus adult data, and in particular for the calculation of cross-brain connectivity between adults and infants. In this final case, it is not clear whether the window size settings should default toward those used for adults or for infants.

There is currently no clear consensus in the field as to how these issues should be addressed. However, it is reasonable to suggest that measures of proximity or (dis-)similarity may be used to index the signal-to-noise ratio yielded by different window size settings. These include the Pearson correlation [13], graph theoretic distance metrics such as geodesic [14] distance, and a recent algorithm that considers the physical location of the nodes not just their value, SimiNet [15], see [16] for a recent review. Here, we address the optimal window size problem for functional connectivity analysis of: (1) infant vs adult neural EEG data, with dyadic interaction as our particular interest case and (2) linear vs nonlinear connectivity metrics.

Two different performance metrics (geodesic distance and SimiNet) were used to determine the most appropriate window size for each analysis.

## II. METHODS

### A. Dataset

The EEG dataset comprised of 37 mother-infant dyads who performed a classic social referencing task. Full details of the task can be found in [11]. In summary, infants were seated in a highchair across a table from their parent (Fig. 1). Each trial consisted of a Demonstration phase, where the adult displayed positive or negative affect toward two novel objects and a Response phase, where the infant was permitted to interact freely between the pair of objects. A total of four pairs of objects were presented four times to the infant in counterbalanced order across trials and participants. Importantly, the window-size calculations reported here were done solely on the EEG data during the Demonstration phase, independent of infant performance in the task, to prevent any selection bias toward desired or expected results.

**Figure 1.**
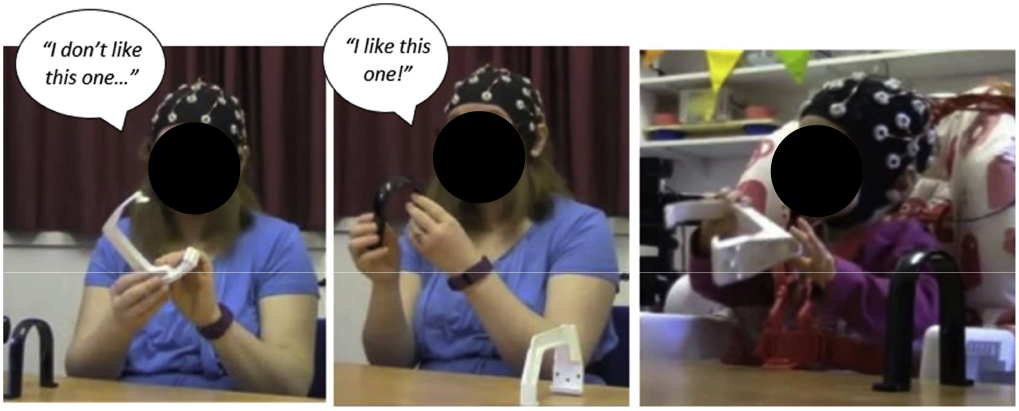
Illustration of experimental setup and task. (Left) Negative object demonstration by adult; (middle) Positive object demonstration by adult; (right) Infant’s interaction with objects. Written informed consent was obtained for the publication of this image.

### B. EEG recordings

Two 32-channel BIOPAC Mobita mobile amplifiers were used with an Easycap electrode system for both infant and adult with a sampling rate of 500Hz. A subset of 16 frontal, central and parietal channels were selected for analysis, consistent with previous analysis performed in [11]. The selected channels were: F_3_, F_z_, F_4_, FC_1_, FC_2_, C_3_, C_z_, C_4_, CP_5_, CP_1_, CP_2_, CP_6_, P_3_, P_z_, P_4_, and PO_z_ (Fig. 2). General EEG preprocessing was performed before calculating the different connectivity metrics: continuous raw data were band pass filtered between 1 and 40 Hz and eye and muscle artifacts were removed with a semiautomatic independent component analysis (ICA) algorithm. Finally, visual inspection of the data was performed to eliminate any residual artifacts before dividing the data into trials. As the number and duration of trials varied across participants, we selected the same number of trials for each dyad, set to 6, and only the first window from the start of each trial was used to avoid statistical bias that may occur when a different number of windows is averaged across trials and participants.

**Figure 2.**
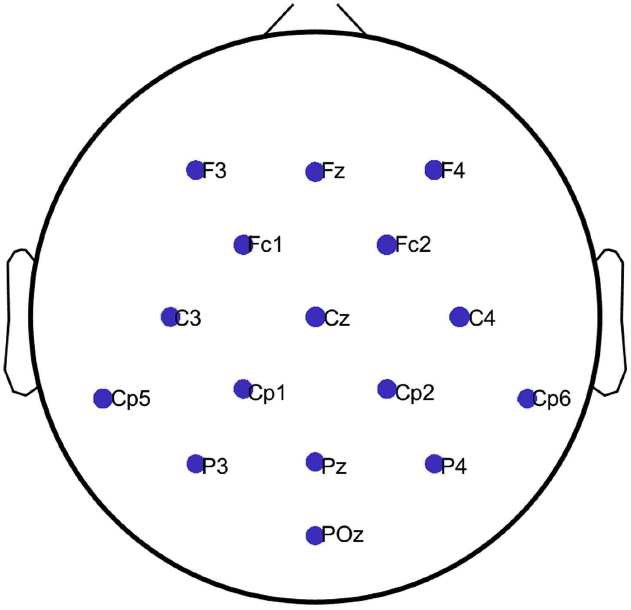
Electrode map of selected channels.

### C. Connectivity metrics

To minimize the effect of volume conduction on our connectivity estimates, we focused on two metrics, the weighted phase lag index (wPLI) [17] and weighted symbolic mutual information (wSMI)[4]. Both of them have been extensively used in EEG-based FC calculations [3], [18], [19].

The PLI is a linear method measuring the extent to which instantaneous phase angle differences between two time series x(t) and y(t) are distributed towards positive or negative parts of the imaginary axis [20]. This asymmetry implies the presence of a consistent, nonzero phase difference (‘lag’) between two time series. The fundamental idea is to disregard phase values centered around 0 phase, as that will be mainly due to volume conduction effects. This also applies at π, 2π and so on. The PLI can be obtained from a time series of phase differences Δ∅(*t*_k_), k=1…N as follows:

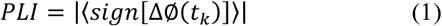

Where sign is the signum function, which will discard a phase difference of 0, π and so on. The PLI ranges between 0, which indicates no coupling and 1, which indicates perfect phase locking. The weighted version of the PLI was introduced to scale the contribution of angular differences based on distance from the real axis, that is, angles closer to zero-lag interaction are regarded as noise affecting global PLI values. The instantaneous phase series was calculated using the analytic signal based on the Hilbert transform. By definition, this should be used for a narrow band, hence we focused on two frequency bands of interest for developmental EEG research, infant ϴ (3-5Hz) and infant α (6-9Hz).

Contrary to wPLI, wSMI is a non-linear metric based on mutual information. It evaluates the extent to which two EEG signals present non-random joint fluctuations, indicating sharing of information[4]. The time series x(t) and y(t) for all EEG channels are first transformed into sequences of discrete symbols 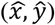. As explained in [4], these symbols are defined by the ordering of *k* time samples separated by a temporal lag τ. Analysis was restricted to a fixed symbol size of *k*=3, implying that symbols comprise of three elements, leading to a 3!=6 possible different symbols and a τ=10 frames, which gives a maximum resolved frequency of 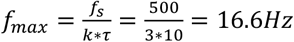. Note, that this means we used a broader frequency band for wSMI than for wPLI. This was done in order to accommodate both infant and adult theta and alpha frequencies. The weights were added in order to discard conjunction of identical and opposite-sign symbols, which indicate spurious correlations due to volume conduction. The wSMI can be calculated as:

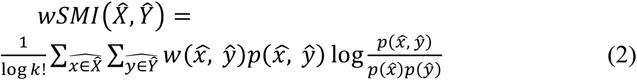

### D. (Dis-)Similarity metrics

Here, two approaches were used to assess signal-to-noise ratio for different window size lengths: **geodesic distance** and the SimiNet graph-based algorithm for **similarity and distance**.

The geodesic distance between two points is the shortest path between them along the manifold as explained in [21] where only one path joins two such points. Translating this concept to FC matrices, *Q*_1_ and *Q*_2_, their geodesic distance can be calculated as:

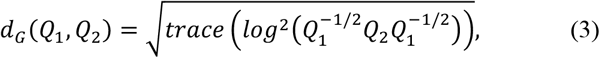

Note that this definition assumes that the matrix *Q*_1_ is invertible. In the cases where this requirement was not met the identity matrix was added as a perturbation matrix to both *Q*_1_ and *Q*_2_ to ensure that all eigenvalues were greater than 0.

In EEG connectivity analysis, the spatial location of the nodes is an important factor to consider when comparing two FC matrices. The intuitive idea behind this is that two networks with the same statistical properties but different regional neural connections should have low similarity [15]. SimiNet is an algorithm that measures the similarity between two graphs taking information from the node and edge properties of the matrices, but also including a spatial constraint related to the physical location of the nodes (i.e. neighbors structure) [15]. The resulting similarity score ranges between 0, indicating completely different graphs and 1, indicating almost identical graphs. This value is based on the sum of a node distance and an edge distance, which is obtained by means of elementary operations. For instance, adding or suppressing nodes until one graph is transformed into the other one. Here we report both values of similarity and distance. Note that the units for geodesic distance and the SimiNet algorithm are not directly comparable.

### E. Bootstrapping

Surrogate datasets (n=1000) were computed per trial through time-point shuffling of the pre-processed EEG dataset for each possible combination and connectivity metric. A non-parametric permutation test (p<0.10) was used as a threshold to suppress likely spurious signals. Afterwards, for each window size, we compared the original (noisier) connectivity metric with its corresponding cleaner (thresholded) version. Accordingly, higher similarity (or lower distance) between the two matrices indicates a more stable extraction of FC measures.

### F. Procedure

Brain connectivity matrices using wPLI and wSMI were calculated for three different window sizes (1s, 1.5s and 2s) and for each member of the dyad (infant, adult, infant-adult) at a trial level. Furthermore, two experimental conditions, positive and negative demonstration phases (see Fig. 1) were evaluated independently. Posteriorly, trial level connectivity matrices were averaged to obtain a single matrix per participant. A similar procedure was performed for the time-domain surrogate values. Finally, the original matrix was compared against the thresholded (bootstrapped) matrix using the different (dis-)similarity metrics as explained above. For each similarity metric, the optimal window size was selected as the one having the highest similarity or lowest (dis-) similarity value. Then, for each dataset and contrast (infant, adult, dyadic infant-adult; for linear wPLI or nonlinear wSMI metrics) we chose the modal window length as the final “winning” outcome.

## III. Results

Recall that two connectivity metrics, wPLI (linear) and wSMI (nonlinear), were used to evaluate optimal window size for infant, adult and infant-adult dyadic datasets. A summary of results can be found in Table 1 and detailed below. Illustrative results are shown in Figures 3 and 4.

**Table 1.** Summary results for optimal window size. Rows are the two connectivity metrics used, wPLI and wSMI. Columns represent evaluations for adult, infant and infant-adult dyadic brain connectivity respectively.

| window size | Adult only | Infant only | Infant-Adult (dyadic) |
| --- | --- | --- | --- |
| wPLI | 2s | 1s | 1s |
| wSMI | 1s | 2s | 1s |

**Figure 3.**
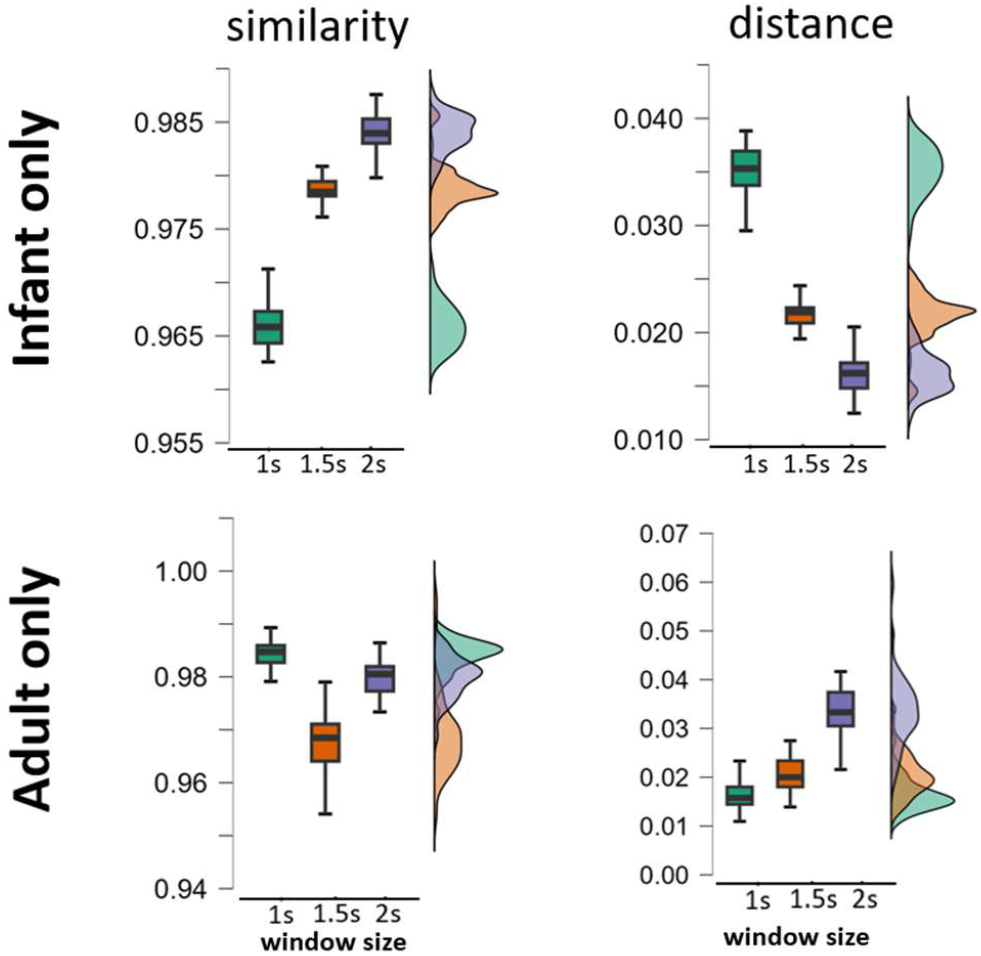
Similarity results for wSMI broadband positive demonstration comparisons. The first row shows the results for the infant only and the second row for the adult only. For illustrative purposes we only show SimiNet results here for Similarity (left column, higher values indicate better signal to noise ratio) and Distance (right column, lower values indicate better signal to noise ratio). Horizontal axes represent the window length, also plotted with different colors. Vertical axes represent the value for each respective metric.

**Figure 4.**
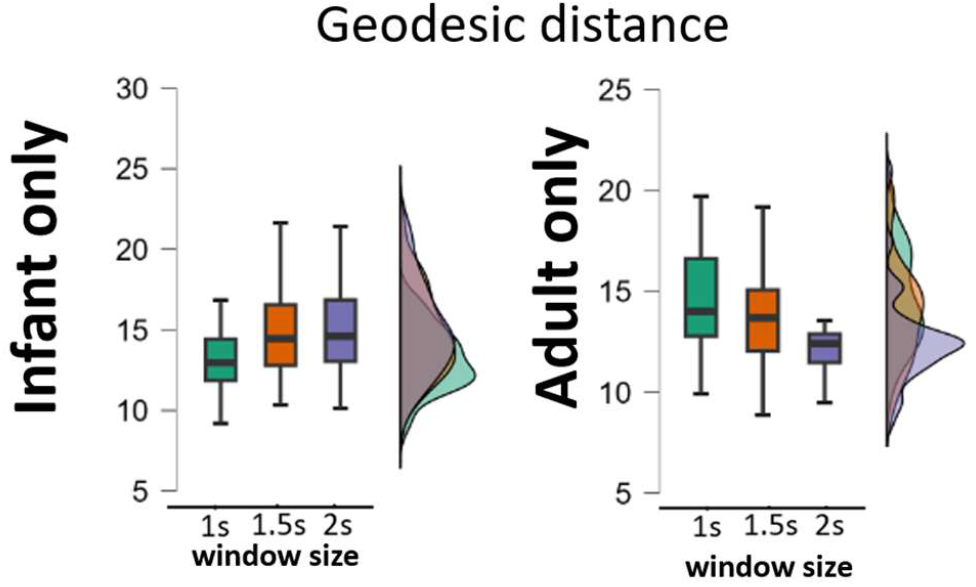
Similarity results for wPLI alpha band negative demonstration condition. For illustrative purposes we only show geodesic distance results for the infant only (left) and adult only (right). Geodesic distance is a measure of dissimilarity, hence the higher the values, the better the signal to noise ratio. are.

### A. Infant vs adult optimal window size: wPLI (linear)

Window size effects on wPLI linear connectivity were assessed for infant ϴ (3-5Hz) and infant α (6-9Hz) frequency bands, for infant, adult and infant-adult dyadic data. For adult EEG data, the longest window (2s length) was the winner, with a more stable pattern observed across similarity metrics, frequency bands and behavioral conditions (7 wins/12 comparisons, or nearly 60% of possible pairwise comparisons).

By contrast, the opposite pattern was observed for infant EEG data where the shortest window (1s length) was the most stable (7/12 or 58.33% of pairwise comparisons). Finally, for infant-mother dyadic EEG, results followed the infant data, with an optimal window size of also the shortest length of 1s (9/12 or 75% of the pairwise comparisons).

### B. Infant vs adult optimal window size: wSMI (non-linear)

Window size effects on wSMI non-linear connectivity (broadband up to 16Hz) were likewise assessed for infant, adult and infant-adult dyadic data. For the adult EEG data, the optimal window size was clearly the shortest length of 1s (5/6, 83% of pairwise comparisons). As for linear measures, the infant data showed the opposite pattern from the adult, where the optimal infant window was the longest length of 2s (5/6, or 83% of pairwise comparisons). Finally, for infant-adult dyadic connectivity, results followed the adult, with an optimal window of 1s (4/6, or 67%).

## IV. Conclusions

Here, we addressed the question of unbiased selection of optimal window size for different linear and non-linear EEG-based functional connectivity metrics, as applied to infant only, adult only and dyadic infant-adult connectivity data. This is a particularly important question for optimizing the detection of true connectivity signals from different developmental datasets, and when studying single versus dyadic (cross-brain connectivity) patterns. Our results indicate that adult and infant connectivity metrics in fact require different window sizes in order to yield stable results, independent of the metric used. We also found that – for the same participant – linear and non-linear metrics require different window size settings. Finally, cross-brain connectivity is not best served by an intermediate window length (i.e., 1.5s), but rather followed either infant or adult settings, depending on the metric used. Another possible interpretation for the results obtained for the dyadic data may be that the optimal window size is independent of the metric used (wPLI or wSMI) and driven by other considerations not captured here.

The optimal time windows observed for infant connectivity data (short for linear metrics, long for nonlinear metrics) are consistent with the idea that the temporal structure of neural interactions may progressively move during development from a regime dominated by oscillatory dynamics to a regime dominated by non-oscillatory dynamics. wPLI should perform more robustly on EEG activity dominated by oscillatory dynamics due to its sensitivity to linear processes. In the case of wSMI, the results observed in infants are also consistent with the idea that longer time windows increase the probability of observing distinct temporal patterns jointly occurring over time, like those characterizing non-linear processes.

Here, we evaluated 1s and 2s as the shortest and the longest time windows respectively. It is thus possible that even more stable FC results may be obtained with window length of <1s and/or >2s, which should be investigated in future. Finally, it should be cautioned that the current results and window size settings should not be overgeneralized, as these may be specific to the task and data type in question. Rather, we have presented a systematic and unbiased approach to evaluating optimal window size settings of infant and adult, single and dyadic EEG data, for linear and non-linear use cases.

## V. Acknowledgments

This research is supported by the RIE2025 Human Potential Programme Prenatal/Early Childhood Grant (H22P0M0002), administered by A*STAR.

## Authors’ background

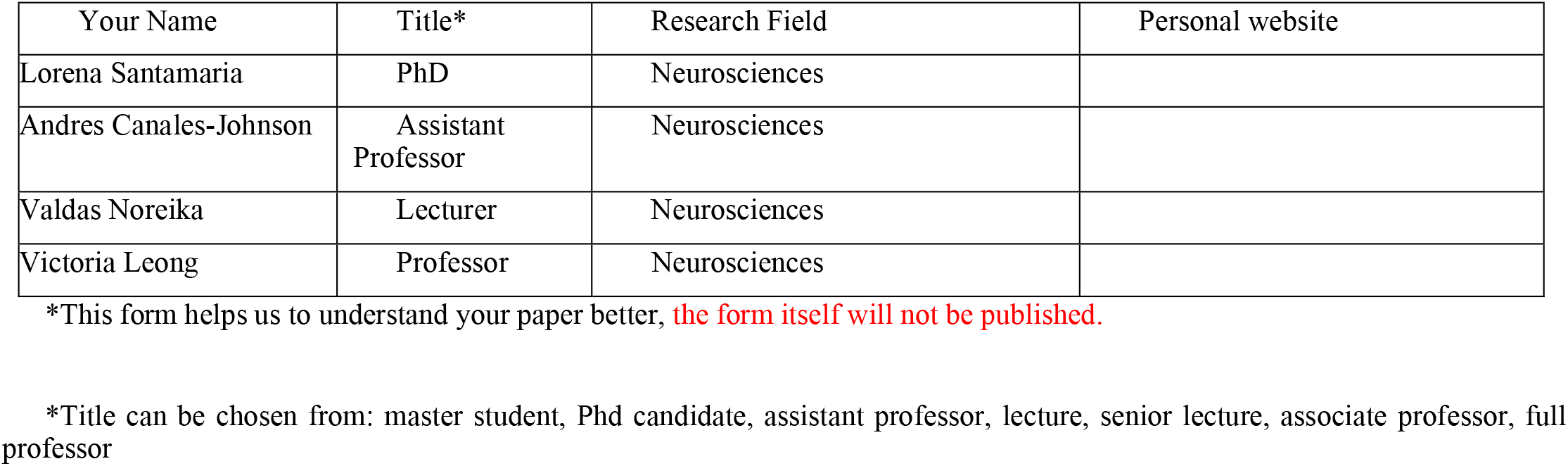

**AUTHORS' BACKGROUND**
| Your Name | Title* | Research Field | Personal website |
| --- | --- | --- | --- |
| Lorena Santamaria | PhD | Neurosciences |  |
| Andres Canales-Johnson | Assistant Professor | Neurosciences |  |
| Valdas Noreika | Lecturer | Neurosciences |  |
| Victoria Leong | Professor | Neurosciences |  |

\*This form helps us to understand your paper better, **the form itself will not be published.**
\*Title can be chosen from: master student, Phd candidate, assistant professor, lecture, senior lecture, associate professor, full professor

